# PACpAInt: a deep learning approach to identify molecular subtypes of pancreatic adenocarcinoma on histology slides

**DOI:** 10.1101/2022.01.04.474951

**Authors:** C. Saillard, F. Delecourt, B. Schmauch, O. Moindrot, M. Svrcek, A. Bardier-Dupas, J-F. Emile, M. Ayadi, V. Rebours, L. de Mestier, P. Hammel, C. Neuzillet, J-B. Bachet, J. Iovanna, N. Dusetti, Y. Blum, M. Richard, Y. Kermezli, V. Paradis, M. Zaslavskiy, P. Courtiol, A. Kamoun, R. Nicolle, J. Cros

**Affiliations:** Owkin France, Paris, France; Université de Paris, Dpt of Pathology, Beaujon Hospital, INSERM U1149, Clichy, France; Dpt of Pathology, Saint-Antoine Hospital - Sorbonne Universités, Paris, France; Dpt of Pathology, Pitié-Salpêtrière Hospital - Sorbonne Universités, Paris, France; Dpt of Pathology, Ambroise Paré Hospital – Université Saint Quentin en Yvelines, Paris, France; Integragen, Paris, France; Université de Paris, Dpt of Pancreatology, Beaujon Hospital, INSERM U1149, Clichy, France; Dpt of Medical oncology, Paul Brousse Hospital, Villejuif, France; Medical oncology, Institut Curie, Paris, France; Dpt of Gastroenterology, Pitié-Salpêtrière Hospital - Sorbonne Universités, Paris, France; Centre de Recherche en Cancérologie de Marseille (CRCM), INSERM U1068, CNRS UMR 7258, Institut Paoli-Calmettes, Aix Marseille Université, Marseille, France; Institut Génétique et Développement de Rennes (IGDR), UMR 6290, CNRS, Université de Rennes 1, Rennes, France; Techniques de l’Ingénierie Médicale et de la Complexité - Informatique, Mathématiques et Applications Grenoble (TIMC-IMAG), CNRS, UMR5525, Université Grenoble-Alpes, Grenoble, France; Centre de Recherche sur l’Inflammation, INSERM U1149, Clichy, France

## Abstract

Pancreatic ductal adenocarcinoma (PAC) is a highly heterogeneous and plastic tumor with different transcriptomic molecular subtypes that hold great prognostic and theranostic values. We developed PACpAInt, a multistep approach using deep learning models to determine tumor cell type and their molecular phenotype on routine histological preparation at a resolution enabling to decipher complete intratumor heterogeneity on a massive scale never achieved before. PACpAInt effectively identified molecular subtypes at the slide level in three validation cohorts and had an independent prognostic value. It identified an interslide heterogeneity within a case in 39% of tumors that impacted survival. Diving at the cell level, PACpAInt identified “pure” classical and basal-like main subtypes as well as an intermediary phenotype and hybrid tumors that co-carried both classical and basal-like phenotypes. These novel artificial intelligence-based subtypes, together with the proportion of basal-like cells within a tumor had a strong prognostic impact.

## Main text

Pancreatic ductal adenocarcinoma (PAC) is predicted to be the second cause of death by cancer in 2030 and its prognosis has seen little improvement in the last decades ^1^. PAC is a very heterogeneous tumor with preeminent stroma and various histological aspects. Omic studies confirmed its intertumor molecular heterogeneity, possibly one of the main factors explaining the failure of most clinical trials. Two transcriptomic subtypes of tumor cells and stroma respectively, were described with major prognostic and theranostic implications ^2–5^. Within the tumor cells, the basal-like subtype is defined by a poorer prognosis linked to early metastases and Folfirinox resistance compared to the classical subtype characterized by a progenitor epithelial phenotype with altered metabolism ^6^. Within the stroma, the activated stroma is enriched in disorganized pro-tumor cancer associated fibroblasts with little extracellular matrix while the inactive stroma is characterized by abundant and dense collagen secreted by more quiescent myofibroblasts. As of today, these subtypes can only be defined by RNA profiling. Tools were proposed to phenotype PAC, either with a binary classification of tumor cells (PurIST basal-like/classical) or with semi-quantitative transcriptomic signatures of tumor (basal-like, classical) and stroma (Activated, Inactivated) subtypes ^5,7^. These approaches are limited by the quantity and quality of the samples (formalin fixation and low cellularity like in biopsies) as well as by the analytical delay and trans-platform reproducibility that restrict its application in clinical trials and in routine care. In addition, tumors may harbor a mixture of several subtypes complicating their interpretation using bulk transcriptomic approaches, thereby limiting their clinical value ^8^. A recent study suggested that tumor cell architecture (i.e formation of glands) could partially predict tumor cell transcriptomic subtypes in primary resected tumors ^9^. This approach requires highly trained pathologists and the analysis of the whole tumor.

Here, we propose PACpAInt, a multi-level AI-based tool using deep learning models to determine molecular subtypes from routine histological preparation (Hematoxylin-eosin-safran (HES staining)), at a resolution enabling to decipher intratumor heterogeneity on a massive scale and providing in addition spatial information of the different cell types and their molecular phenotype (Fig.1a). The model was trained and validated on multicentric cohorts using 1796 slides (602 patients) with corresponding transcriptome from the same lesion (cohorts are described in Extended Fig 1). Deep learning models were trained on a discovery cohort (DISC cohort) composed of 424 whole-slide histological images from 202 PAC resected in 3 centers (mean number of slides/case = 2) in which the neoplastic areas were annotated by two pathologists. These models were externally validated in 3 cohorts of resected PAC (BJN_U (n=148), BJN_M (n=97), TCGA (n=126)) and one cohort of liver fine-needle biopsies (Liver_FNB, n=25) from PAC metastatic cases (Extended Data Fig.1). For BJN_M and FNB, RNASeq was specifically performed on the same tissue area as the one processed by the algorithms (i.e. “spatially matched”).

**Figure 1:**
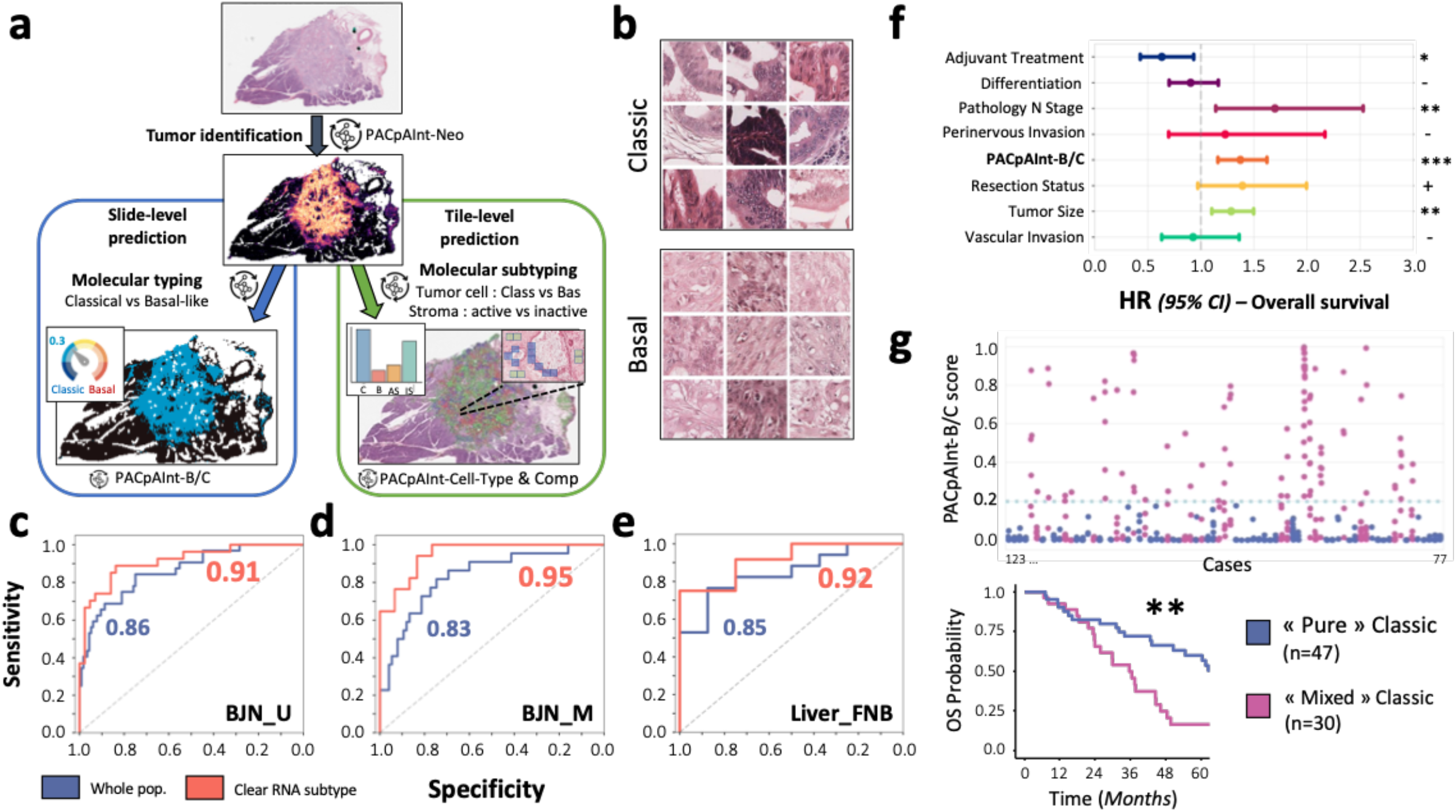
PACpAInt identification of PDAC molecular tumor subtypes. a) Simplified workflow of the study: a first model is applied to find the tumor (PACpAInt-Neo) followed by a second model predicting either the global tumor cell molecular type (classical vs basal-like) at the slide level (PACpAInt-B/C) or predicting at the tile level (small square 112μm wide) the nature of the cells (tumor or stroma - PACpAInt-cell type) and their molecular subtype (classical vs basal-like for tumor cells and active vs inactive for stroma) (PACpAInt-Comp), b) Representative tiles identified as classical or basal-like by PACpAInt-B/C in the validation BJN_U cohort, c/d/e) Performance of PACpAInt-B/C to identify molecular subtypes in validation cohorts (BJN_U unmatched (surgical specimens), i.e slide analysed and tissue used for RNAseq are not spatially matched; BJN_M matched (surgical specimens), i.e slide analysed and tissue used for RNAseq are spatially matched; Liver_FNB (EUS fine needle biopsies), g) Multivariate analyses of clinical/pathological factors and PACpAInt-B/C demonstrating an independent prognostic value of the later on overall survival, f) Application of PACpAInt-B/C to all the tumor slides (n=660) of 77 cases defined as classical by RNAseq on a single sampling. Top panel: The PACpAInt-B/C score estimating the “basalness” of each slide is represented on the Y axis while patients (1 to 77) are lined along the X axis. Each spot represents a slide. Cases with all their slides showing a low PACpAInt score (<0.2) were called “pure” classical compared to more heterogeneous tumors called “mixed” classical. Bottom panel: Kaplan Meyer analysis of overall survival comparing “pure” and “mixed” classical tumors. ***: p < 0.001; **: p < 0.01; *: p < 0.05; +: p < 0.1; -: p > 0.1.

A first model (PACpAInt-Neo) was developed to predict neoplastic areas and successfully detected those regions when run on two independent validation cohorts (AUC: BJN_U=0.99, TCGA=0.98) (Extended Data Fig.2). The second model (PACpAInt-B/C) was trained on the DISC cohort using only predicted neoplastic areas to determine the PurIST-RNA basal-like/classical (B/C) subtypes. Despite the high histological diversity of PAC, PACpAInt-B/C identified a set of morphological features specific to both basal-like and classical subtypes (Fig 1.b). For the task of predicting RNA subtypes basal-like/classical, PACpAInt-B/C was successfully validated in 2 spatially unmatched validation cohorts, with AUCs of 0.86 [0.79 - 0.94] and 0.81 [0.71 - 0.90] in the BJN_U and TCGA cohorts respectively (Fig.1c & Extended Data Fig.3a/b). Comparable performance was achieved on a third validation cohort with spatially matched histological and molecular areas (BJN_M, AUC=0.83 [0.73-0.93])(Fig.1d).

Given the previously described intratumor heterogeneity, we also restricted the analysis to the 50% of cases that had the clearest, unambiguous transcriptomic subtype and showed that the performance improved substantially (AUC of 0.91 [0.84 - 0.98] and 0.88 [0.79 - 0.98] in the BJN_U and TCGA cohorts respectively (Fig.1c and Extended Data Fig.3a) ^8,10^. This was particularly significant within the spatially matched validation cohort BJN_M (AUC=0.95 [0.90 - 1.0]) highlighting the limitations of a binary classification in highly heterogeneous tumors (Fig.1d). Because most patients are diagnosed at the metastatic stage on liver biopsies, we validated PACpAInt-B/C on 25 fine needle liver biopsies with matched RNAseq data (FNB cohort). Performance remained as good as for surgical specimens (AUC=0.85 [0.69 - 1.0]), and similarly improved in cases with a clear molecular subtype (AUC=0.92 [0.77 - 1.0], (Fig.1e). We included PACpAInt-B/C in a multivariate survival analysis in the pooled BJN_U and BJN_M cohorts. PACpAInt-B/C predictions had a strong independent prognostic value on both OS (HR=1.37 [1.16 - 1.62] p<0.001) and DFS (HR=1.27 [1.08 - 1.49] p=0.003), contrary to the PurIST-RNA classification which was not associated with OS (Fig.1f, Extended data Fig.3c-3d and Extended Table 1-2).

It has been previously shown that tumor cells may harbor distinct morphology from slide to slide within a case ^9^. This is particularly meaningful in tumors of the classical subtype which may harbor basal-like areas that could impact patient prognosis. In order to assess the impact of minor basal-like areas, we ran PACpAInt-B/C on all available tumor slides (mean nb of slides/cases = 9) from the RNA-defined classical cases in BJN_M validation cohort (n=77/97) and compared predictions across slides (Fig.1g). 30 (39%) cases had at least one slide predicted as basal-like, suggesting an important morphological and molecular heterogeneity within those tumors. DFS and OS of these heterogeneous cases were shorter (median survival of 15 vs 35 months, p=0.08 and of 36 vs 64 months, p=0.002 respectively), highlighting the clinical impact of tumor heterogeneity (Fig.1g).

We further explored this intratumor heterogeneity by decomposing the whole slide image into small squares measuring 112 micron-wide, called tiles and analyzing each of them individually (mean nb of tiles / slide = 23,306, Extended Data 1.c). PACpAInt-CellType model was trained on pathological annotations to differentiate tumor cells from stroma within the neoplastic area at the tile level. The model reached an AUC of 0.99 in the two validation cohorts BJN_U and TCGA (Extended Data Fig.4a/b). PACpAInt-CellType was further validated on a subset of BJN_M (n=50) for which the tumor cell/stroma ratio was digitally computed using a tumor specific pan-cytokeratin immunohistochemistry (Pearson’s R=0.72, p<0.001) (Extended Data Fig.4c). Using this model predictions on a pooled cohort (DISC, BJN_M and BJN_U, n=451), we confirmed that a high amount of stroma was independently associated with better prognosis (HR=0.86 [0.76-0.96], p=0.01 and HR=0.87 [0.77 - 0.98], p=0.02 for DFS and OS, respectively) (Extended Data Fig.4d and Extended Table 3), as formerly reported ^11–13^.

A final model PACpAInt-Comp was developed to recognize at the tile-level stroma molecular subtypes (active/inactive) in addition to tumor cell subtypes (basal-like/classical) (Extended Data Fig. 1.c) ^5^.

The correlation of PACpAInt-Comp with RNA tumor and stroma signatures was highly significant in the BJN_U validation cohort and substantially improved in the spatially matched validation cohort BJN_M (Fig.2a). Concordance between tile level predictions of the model (basal-like/classical) on tumor cell tiles and active/inactive on stroma tiles) and tile scoring by two expert pathologists in PAC was excellent (concordance = 100% and 99.2% for tumor components and 99.2% and 99.4% for stroma components). In addition, tiles predicted to have a basal-like and classical phenotype according to PACpAInt-COMP were tiles containing tumor cells (according to PACpAInt-CellType). Similarly, tiles predictive of containing active or inactive stroma (stromaActive or StromaActiv component high score respectively) according to PACpAInt-COMP were tiles containing stroma (at not tumor cells) according to PACpAInt-CellType (Extended Data Fig.5a). PACpAInt-Comp was further evaluated on slides stained with GATA6/Claudin18 and KRT17 antibodies, three established specific markers for classical and basal-like phenotypes respectively ^14,15^. PACpAInt-Comp was able to discriminate between GATA6+/Claudin18+ versus KRT17+ areas, with AUCs of 0.87 and 0.75 for basal-like and classical tumor scores, respectively (Fig 2.b). Using PACpAInt-Comp predictions, we also highlighted a strong association between active stroma and basal-like rather than classical phenotypes (Extended data Fig.5b) as previously reported ^5^. In a multivariate Cox model, the use of the PACpAInt-Comp scores significantly improved the prognosis prediction with respect to clinico-molecular data alone (+4 c-index, p=0.007 and +3 c-index, p=0.008 for OS and DFS respectively) (Extended Table 4).

**Figure 2:**
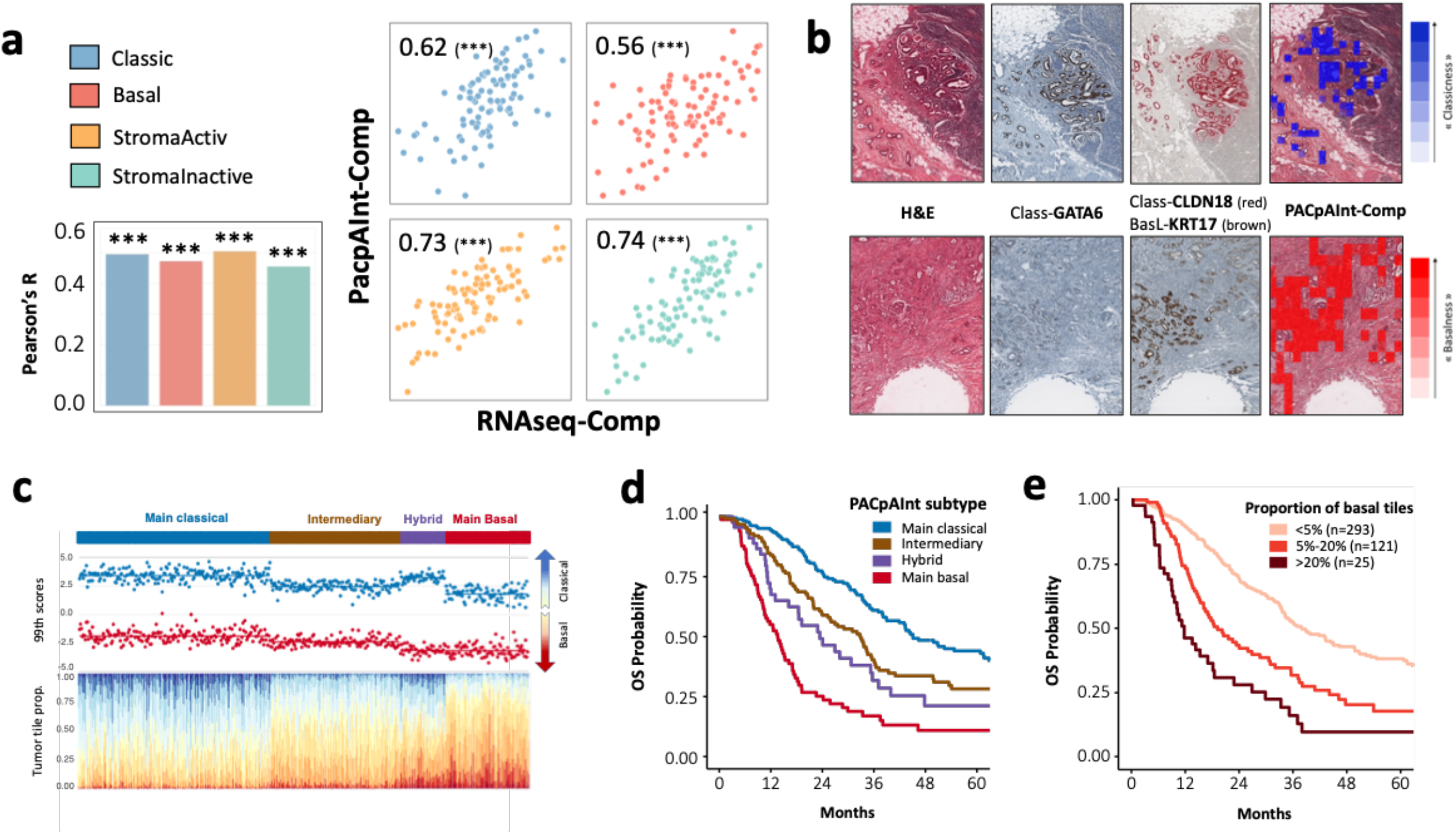
PACpAInt identification of PDAC molecular components to decipher intratumor heterogeneity. a) Correlation at the slide level between the tumor and stromal components defined by RNAseq or PACpAInt-Comp on the BJN_U unmatched (left panel) or BJN_M matched (right panel) validation cohorts, b) exemple at the tile level of areas identified as classical or basal-like by PACpAInt-Comp and stained by immunohistochemistry with classical (GATA6/Claudin18) or basal-like (KRT17) markers, c) Patient distribution in four subtypes: main-classical, intermediary, hybrid and main-basal-like. For each column-wise patient is first shown the 99th percentile basal-like and classical scores and second the proportion of tumor tiles for different levels of basal-like and classical phenotype. d) Kaplan Meyer analysis of overall survival comparing main classical, intermediate, hybrid and main basal-like tumor, e) Kaplan Meyer analysis of overall survival comparing tumors with less than 5%, 5 to 20% and more than 20% of their tumor tiles being identified as basal-like.

We further assessed the impact of intratumor heterogeneity, using PACpAInt-Comp to spatially phenotype a total of 6.3 million tumor tiles encompassing 451 patients of the pooled cohort (DISC, BJN_U and BJN_M).

The tile by tile analysis of the basal-like and classical PACpAInt-Comp scores showed that only 60% of tumors had an unambiguous main subtype (classical 41%, basal-like 19%). The remaining could be divided into an infrequent hybrid subtype (10%) defined by the coexistence of both clearly differentiable basal-like and classical tumor cells, and an intermediary subtype (30%) for which most tumor cells could not be clearly assigned to any of the two subtypes (Fig.2c). The latter two subtypes had an intermediate prognosis (Fig.2d and Extended Data Fig.5c). PACpAInt subtyping of the pooled cohort with 451 cases revealed that 71% of tumors present a detectable fraction of clearly basal-like identified tumor cells, confirming that most PDAC tumors contain basal-like cells as suggested by previous single-cell analyses on 12 cases^8,16^. The overall proportion of basal-like cells was prognostic, with worsen prognosis starting at 5% of clearly basal-like identified tumor cells (Fig. 2e and Extended Data Fig. 5d), and was independently associated with OS and DFS in a multivariate analysis (Extended Data Fig. 5e and Extended Table 5).

## Discussion

With a global consensus on PAC molecular subtypes finally emerging and early results suggesting their potential predictive value in addition to a strong prognostic value, the need for efficient and reliable tumor subtyping is greater than ever ^6^. In this study we developed PACpAInt, the first AI-based tool able to predict on routine pathology slides PAC molecular subtypes of both tumor and stromal cells. The originality of our approach is to rely on an interpretable deep learning design, translating molecular signatures defined on whole tumors into morphology-based spatialized cell phenotyping for comprehensive intratumor heterogeneity analyses.

Our training cohort included slides from different centers over a long period of time with different staining protocols ensuring a wide variability in stainings in order to build robust models. The very good performance on the validation cohorts from a fourth center and the multicentric TCGA cohort supports the robustness of the models. Most importantly, the model performed well on liver biopsies, the most common diagnostic sample for PAC diagnosis. The binary classification of tumor cells could help decide the treatment instantly, without the lengthy and costly RNAseq analysis, allowing its use in clinical trials to stratify patients. In addition, it could detect the remaining tumor cells after neoadjuvant treatment, paving the way for a standardized regression score that could also be used in trials to adjust the adjuvant therapy. PACpAInt allowed us to assess the intratumor heterogeneity at a level never explored before. Few studies performed multi-areas RNAseq, each on a small number of cases, suggesting that the two main subtypes may be present in a single tumor ^17,18^. Our results provide the first clear picture of PAC intratumor heterogeneity showing that almost a third of the tumors are likely halfway between the classical and the basal-like subtypes. This is of major interest as several epigenetic drugs are being developed to try to reprogram PAC cells. Our data showed that a minor basal-like component, that would be ignored by binary classifications, has a strong prognostic implication. Finally, this study also demonstrates that the stromal compartment can be rapidly subtyped, paving the way for patient stratification in drug targeting trials.

In conclusion, while demonstrating the value of histology-based deep learning models for tumor subtyping in PAC, these results also show the limit of molecular-based subtyping in highly heterogeneous samples. With the expansion of digital pathology, remote AI-based PAC subtyping could be deployed worldwide, finally opening the way for patient stratification based on powerful molecular criteria.

## Extended Methods

### Datasets description

The discovery set used to develop our models is a multicentric cohort of 202 patients treated in Saint-Antoine University Hospital, Pitié-Salpêtrière University Hospital or Ambroise Paré University Hospital, between september 1996 and december 2010. At least two hematoxylin-eosin +/− safran (HES) slides from surgical specimens were available for each patient, corresponding to a total of 424 slides. BJN_U, BJN_M are two independent validation cohorts of patients treated at Beaujon University Hospital between september 1996 and january 2014. BJN_U consists of 304 HES slides of surgical resection specimens from 148 patients. For the BJN_M cohort, all slides of the tumor specimens were digitized, corresponding to a total of 909 HES slides for 100 patients. FNB is a third independent validation cohort of endoscopy ultrasound fine needle biopsies from liver metastasis of 25 patients (one biopsy per patient) treated at Beaujon University Hospital between 2013 and 2020. TCGA is a multicentric independent validation cohort of 134 hematoxylin-eosin (H&E) slides (126 cases) from a public dataset of the TCGA database ^19^. Inclusion criteria for all cohorts were as follows: unequivocal diagnosis of PAC, available histological slides of formalin-fixed, paraffin-embedded material, available follow-up and molecular information, absence of metastasis at diagnosis. This led to the exclusion of 34 slides from the TCGA that had either no tumor cell on the slide, or were from frozen examinations.

### Transcriptome profiling and molecular subtyping

The discovery cohort corresponds to 202 resected tumors from the Puleo et al study which were profiled using U219 Affymetrix microarrays. For the BJN_U cohort, RNA was extracted from a 0.8mm diameter core sampled from a tumor-enriched zone. In most cases, the RNA was not extracted from the same block that was used to generate the HES slides. For the BJN_M and FNB series, RNA was extracted after manual microdissection to remove contaminating normal liver or pancreatic tissue. For these cohorts, the RNA was extracted from exactly the same area as the analysed HES. In addition, for the BJN_M cohort, all the other tumor slides were also analyzed by PACpAInt. For the BJN cohorts, DNA/RNA was extracted using the ALLPrep FFPE tissue kit (Qiagen, Venlo, The Netherlands) following the manufacturer’s instructions and sequenced using 3’ RNAseq (Lexogene Quantseq 3’). RNAseq reads were mapped using STAR and genes were quantified using FeatureCount. Gene-counts were Upper Quartile-normalized and logged. PurIST was applied to both microarray and RNAseq profiles resulting in a class label for each sample. The tumor and stroma components were applied to both microarray and RNAseq profiles resulting in a continuous score for each component in each sample as previously reported ^20^. For each dataset, the difference between the scaled basal-like and classical component scores were computed and samples that had a difference above the median were considered to have a clear RNA subtype.

### Preprocessing of whole-slide images

The application of deep-learning algorithms to histological data is a challenging problem, particularly due to the high dimensionality of the data (up to 100,000 × 100,000 pixels for a single whole-slide image) and the small size of available datasets. Therefore a preprocessing pipeline composed of multiple steps was used to reduce dimensionality and clean the data. First step consists in detecting the tissue on the WSI: a U-Net neural network is used to segment part of the image that contains matter, and discard artifacts such as blur, pen marker etc., as well as the background ^21^. Second step consists in tiling the slide into smaller images, called “tiles”, of 112 × 112 μm (224 × 224 pixels). At least 20% of the tile must have been detected as foreground by the U-Net model to be considered as a tile of matter. During training, a maximum of 8000 such tiles are then uniformly sampled from each slide. Final step consists of extracting features from each tile; 2,048 relevant features are extracted using a self-supervised model, MoCo v2, using the approach proposed by Dehaene et al ^22^. At the end of this preprocessing pipeline, each slide is represented by a matrix of size (n_tiles_, 2048).

### Neoplastic and cell type Prediction

PACpAInt neoplastic prediction model (PACpAInt-Neo) was trained at the tile level, based on neoplastic annotations of 433 slides from the discovery cohort provided by two expert pathologists, which corresponds to a total of 9,886,596 tiles. WSI preprocessing described in section “Preprocessing of whole-slide images” was used to obtain 2,048 features for each tile. PACpAInt-Neo consists of a multi-layer perceptron with a single layer of 128 hidden neurons, followed by ReLU activation. The model was validated on regions annotated by two pathologists of slides of the cohort BJN_U and TCGA (10 slides for each cohort). PACpAInt cell type prediction model (PACpAInt-Cell type) has the same architecture as PACpAInt-Neo, and was trained on annotations of 81 slides from the discovery cohort, which corresponds to a total of 66920 tiles. Likewise, it was validated on regions of 10 slides of BJN_U and TCGA_PAAD annotated by two pathologists.

### Molecular Prediction

PACpAInt-B/C and PACpAInt-Comp are two deep learning models that were trained on the discovery cohort to predict respectively PurIST basal-like classification and the molecular components classical, Basal-like, StromaActiv, StromaInactive. The two models use the same WSI preprocessing pipeline described in section “Preprocessing whole-slide images”. PACpAInt-Neo was further applied on the tile features in order to select only tiles in neoplastic regions (i.e. with a neoplastic prediction score larger than 0.5). PACpAInt-B/C architecture is similar to the one proposed by Ilse et al.: A linear layer with 128 neurons is applied to the embedding followed by a Gated Attention layer with 128 hidden neurons ^23^. We then apply a MLP with 128 and 64 hidden neurons and ReLU activations to the results. A final Sigmoid activation is applied to the output to obtain a score between 0 and 1, which represents the probability of the slide to be basal-like or classical. PACpAInt-B/C was trained with the binary cross entropy as loss function. PACpAInt-Comp was inspired by the WELDON algorithm: A set of one dimensional embeddings is computed for the tile features using a multi-layer perceptron (MLP) with 128 hidden neurons followed by 4 neurons and ReLU activation ^24^. For each channel output of the MLP, we select R=100 top and bottom scores and average them, so that the model’s output is a vector of size 4, corresponding to the predicted values of each molecular component. PACpAInt-Comp was trained with the mean squared error as loss function.

### Spatial Validation

To validate locally the accuracy of PACpAInt-Comp to predict classical and basal-like, GATA6 and KRT17 IHCs were performed on 12 slides of BJN_M. Tile scores for classical and basal-like components were analyzed in regions defined as being basal-like/classical by the IHCs. Two expert pathologists also analysed tiles predicted to be classical or basal-like (n=500) and tiles predicted to be stroma active or inactive (n=500), blinded to scores associated with each tile.

### Performance assessment and statistical methods

The area under the receiver operating characteristic curve (AUC) was used to quantify the capability of the model to distinguish classical from basal-like tumors, as assessed by the PurIST method. The same metric was used to assess the performance of PACpAInt-Neo to distinguish normal from neoplastic regions, and of PACpAInt-Cell-Type to distinguish stroma from epithelial tumor cells. Delong’s method was used to compute confidence intervals at 95% confidence level ^25^. Pearson’s correlation was used to assess the performance of the PACpAInt-Comp model to predict the molecular components. Survival analyses were performed with uni- and multivariate Cox proportional hazards models implemented in the lifelines package of Python ^26^. Log-rank tests were used to compare survival distributions between population subgroups. We used survcomp R package to compare c-indexes ^27^. All tests were twotailed, and P values < 0.05 were considered statistically significant.

### Intratumor heterogeneity subtypes

The 99th percentile of the basal-like and classical component scores defined by PACpAInt-Comp for each patient, using all available slides per patient, were computed. The difference between 99th basal-like and classical was then use to distinguish two groups of tumors using a mixture of gaussians, a high difference group of tumors was considered as well-identified either as basal-like or classical, and a low difference group of tumors with ambiguous basal-like/classical subtyping. The maximum of the basal-like or classical components was then used to separate further the low difference group, high maximum group with low difference between extremes were termed hybrid given both high classical and basal-like phenotype, while the low maximum group with low difference between extremes was termed intermediary as no tumor cell tile reached a high level of either basal-like or classical phenotype.

### Clinical variables used in the multivariate analysis

Clinical variables considered for multivariate analysis were common variables known to be associated with PAC prognosis: pN stage, differentiation, perineural invasion, resection status, tumor size, vascular invasion, adjuvant treatment yes/no.

### Ethical compliance

The project (ref 2020-013) was reviewed and approved by the “Comite d’Evaluation de l’Ethique des projets de Recherche Biomedicale (CEERB) Paris Nord” (Institutional Review Board-IRB 00006477-of HUPNVS, Paris 7 University, AP-HP).

---

**Extended data 1:**
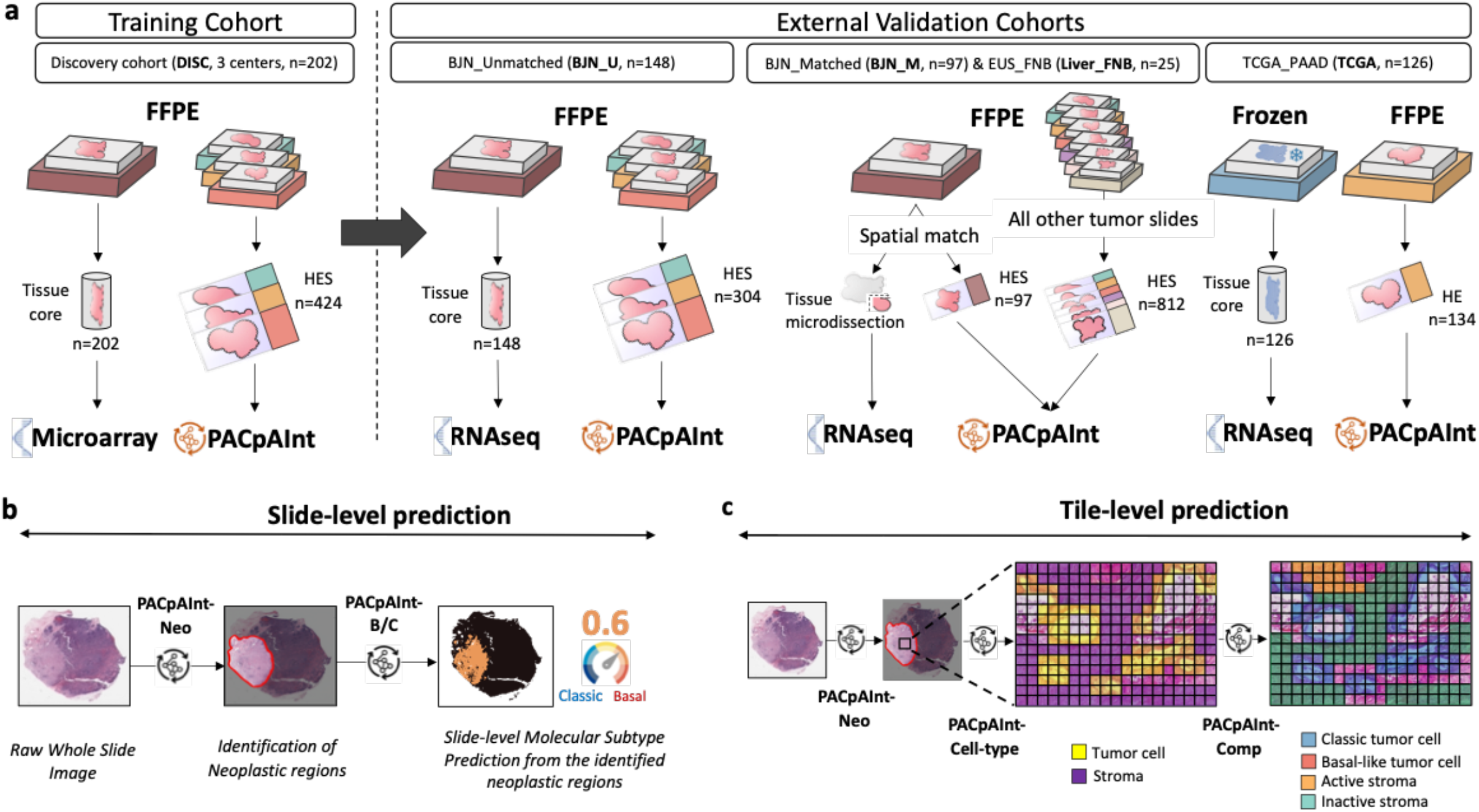
Detailed flow chart of the study. a) Description of the cohorts. Discovery cohort was composed of 202 patients (surgical specimens) from 3 centers. A tissue carrot (diameter 600m) was taken from a block for RNA profiling. HES slides (at least 2/tumor) were digitized for PACpAInt analysis. In most cases the tissue carrot and the HES did not come from the same block. The workflow was similar in the first validation cohort BJN_U unmatched (surgical specimens). For the 2 next validation cohorts (BJN-M matched (surgical specimens) and EUS_Liver (liver metastases, fine needle biopsies)), the same block was used for RNA extraction after microdissection of neoplastic area and to generate the HES slide that was digitized and analyzed with PACpAInt. In addition, in the BJN_M matched cohort, all the remaining tumor slides were also digitized for PACpAInt analysis. Finally, in the TCGA_PAAD validation cohort (surgical specimens), in contrast to all the other cohorts, the RNA was extracted from frozen material, not formalin-fixed paraffin-embedded. Similarly to the discovery cohort, the tissue analyzed by RNAseq was not spatially matched with the digitized slides. b) Slide-level prediction. At the whole slide level (global classification of the whole slide), the multistep PACpAInt model first recognizes neoplastic areas (PACpAInt-Neo module), then assesses the basal-like of classical status (PACpAInt-B/C module). c) Tile-level prediction. In this setting, all the tiles (small square, 112um wide) are analyzed and reported individually. The multistep PACpAInt model first recognizes neoplastic tile (PACpAInt-Neo module), then recognize tumor cell and stroma (PACpAInt-Cell type module) then assesses the molecular components (classical vs basal-like for tumor cell tiles and activated vs inactivated for stroma tiles) (PACpAInt-Comp module) allowing the deep study of intratumor heterogeneity.

**Extended data 2:**
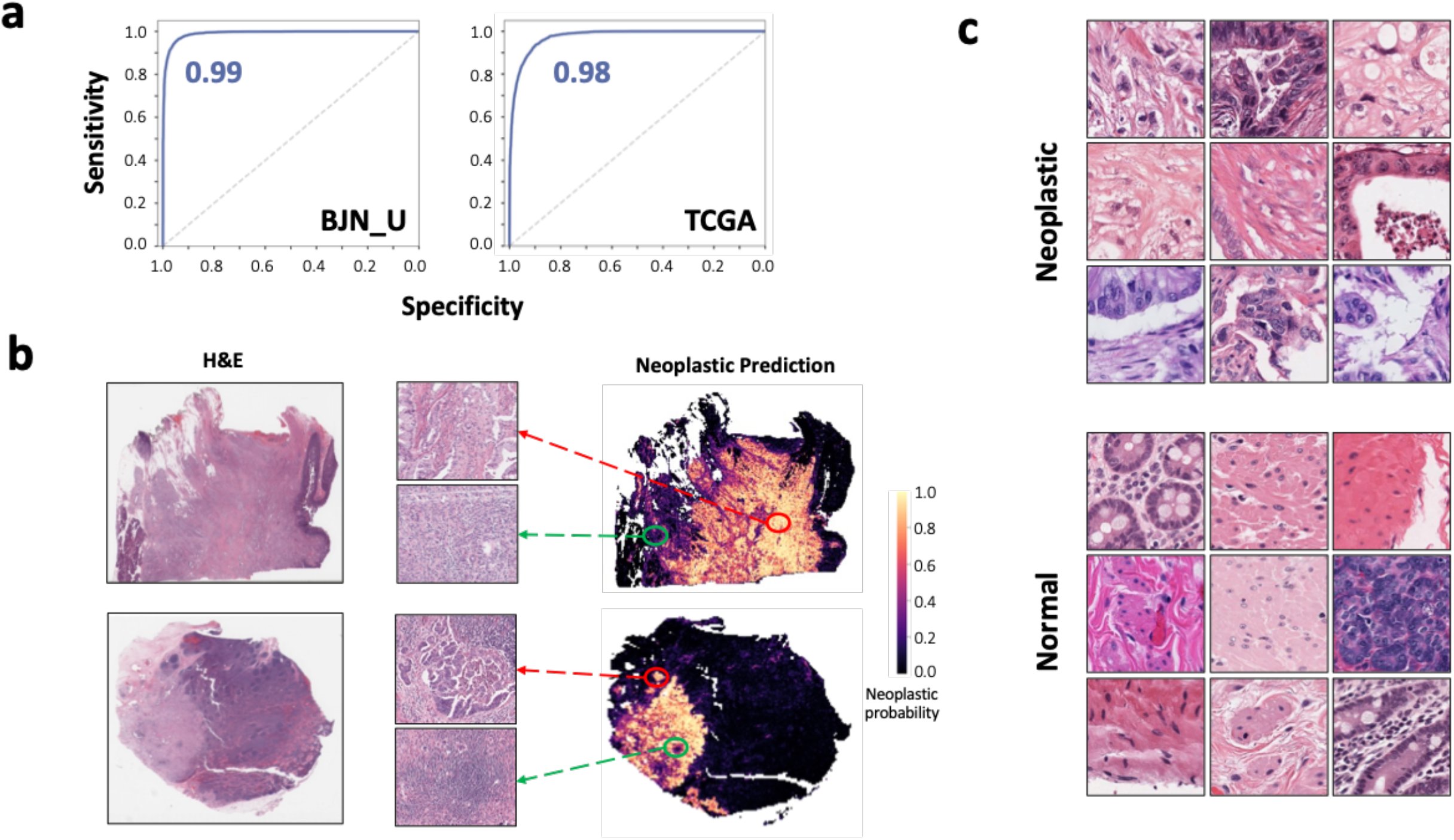
Identification of neoplastic areas by PACpAInt-Neo. a) Performance of PACpAInt to identify neoplastic area in the BJN_U and TCGA validation cohorts, b) Example on 2 cases of neoplastic areas identification with H&E (left), PACpAInt-Neo segmentation (right) and zooms (center) of neoplastic (yellow) and non-neoplastic (green) areas, c) Representative tiles identified as neoplastic and non-neoplastic by PACpAInt-Neo in the TCGA validation cohort.

**Extended data 3:**
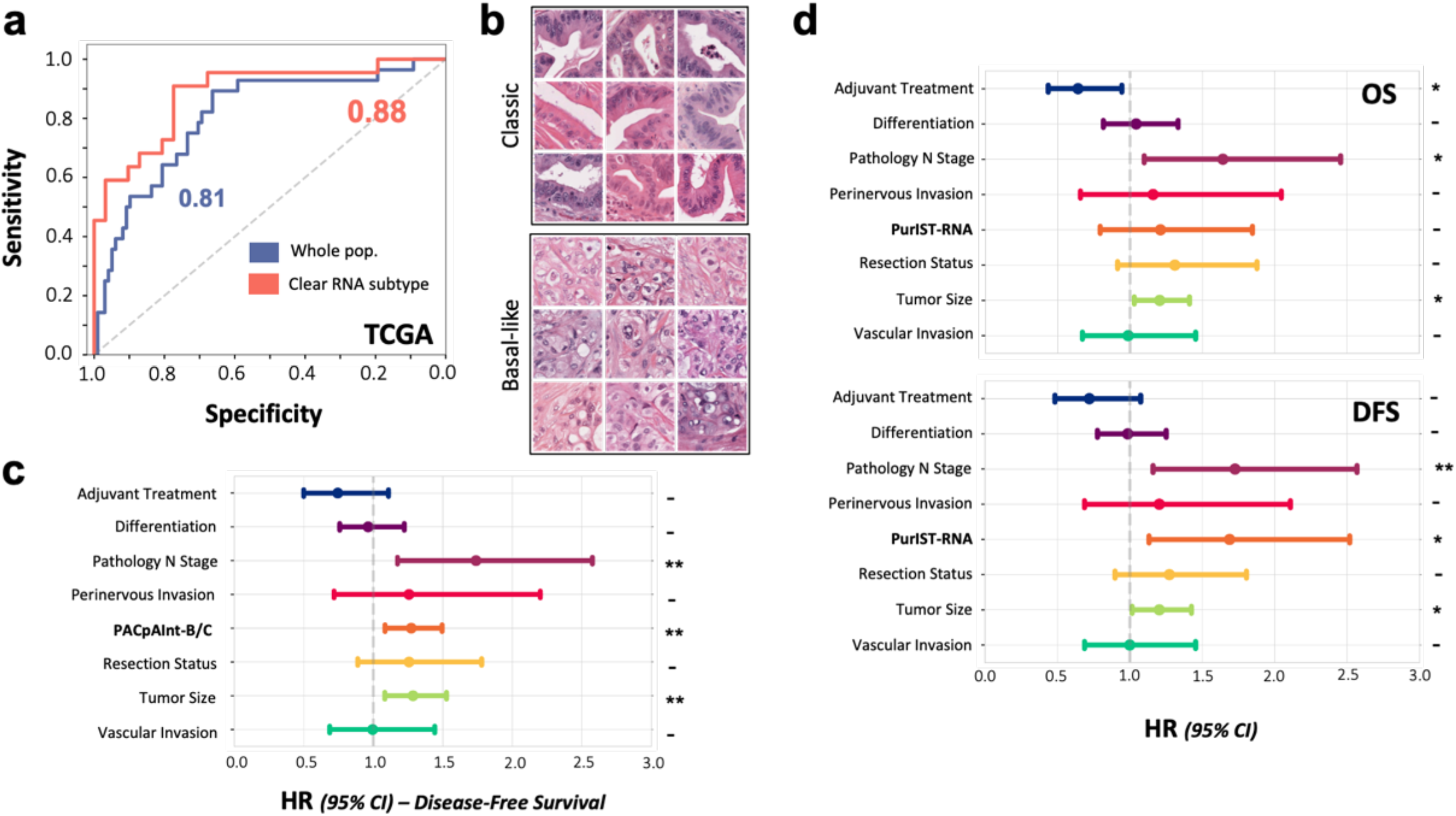
Identification of the molecular subtypes PACpAInt-B/C. a) Performance of PACpAInt-B/C to identify molecular subtypes at the whole slide level area in the TCGA validation cohort using the whole cohort or only cases with an unambiguous RNA subtype (clear subtype), b) Representative tiles identified as classical or basal-like by PACpAInt-B/C in the TCGA validation cohort, c) Multivariate analyses of clinical/pathological factors and PACpAInt-B/C on disease free survival in the BJN_U+M validation cohorts, d) RNA-defined molecular subtype (PurIST-RNA) on overall survival (top panel) and disease free survival (bottom panel) in the BJN_U+M validation cohorts. ***: p < 0.001; **: p < 0.01; *: p < 0.05; +: p < 0.1; -: p > 0.1.

**Extended data 4:**
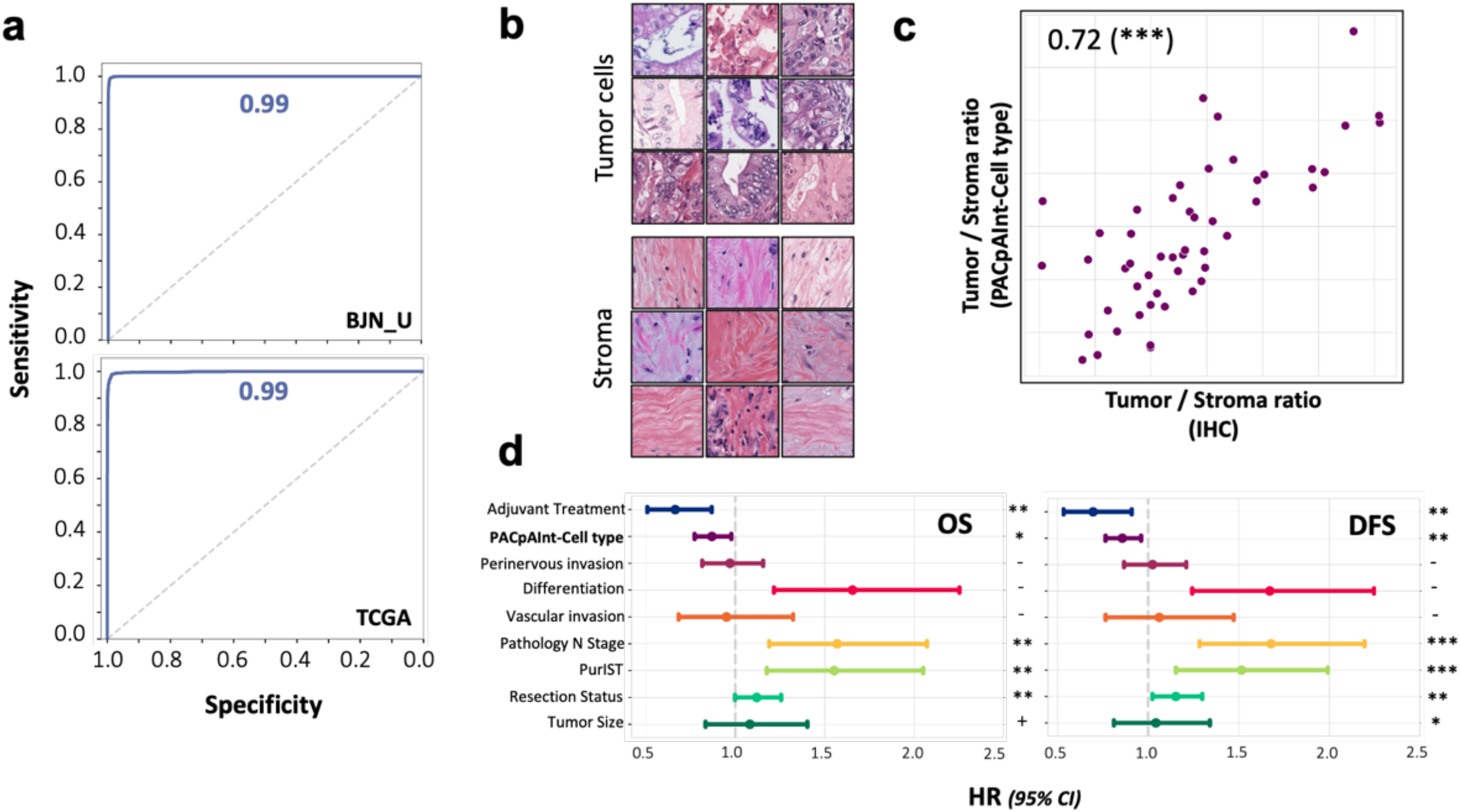
Identification of tumor and stroma cells by PACpAInt-Cell type. a) Performance of PACpAInt-Cell type to identify tumor and stroma cells in the BJN_U (top) and TCGA (bottom) validation cohorts, b) Representative tiles identified as tumor or stroma by PACpAInt-Cell type in the TCGA validation cohort, c) correlation between the tumor cell/stroma ratio computed by PACpAInt-Cell type or with a pan-cytokeratin immunohistochemistry (ratio stained area/total tumor area), d) Multivariate analyses of clinical/pathological factors and PACpAInt-cell type computed tumor/stroma ratio on disease free (left) and overall (right) survival in the pooled DISC cohort and BJN_U+M validation cohorts. ***: p < 0.001; **: p < 0.01; *: p < 0.05; +: p < 0.1; -: p > 0.1.

**Extended data 5:**
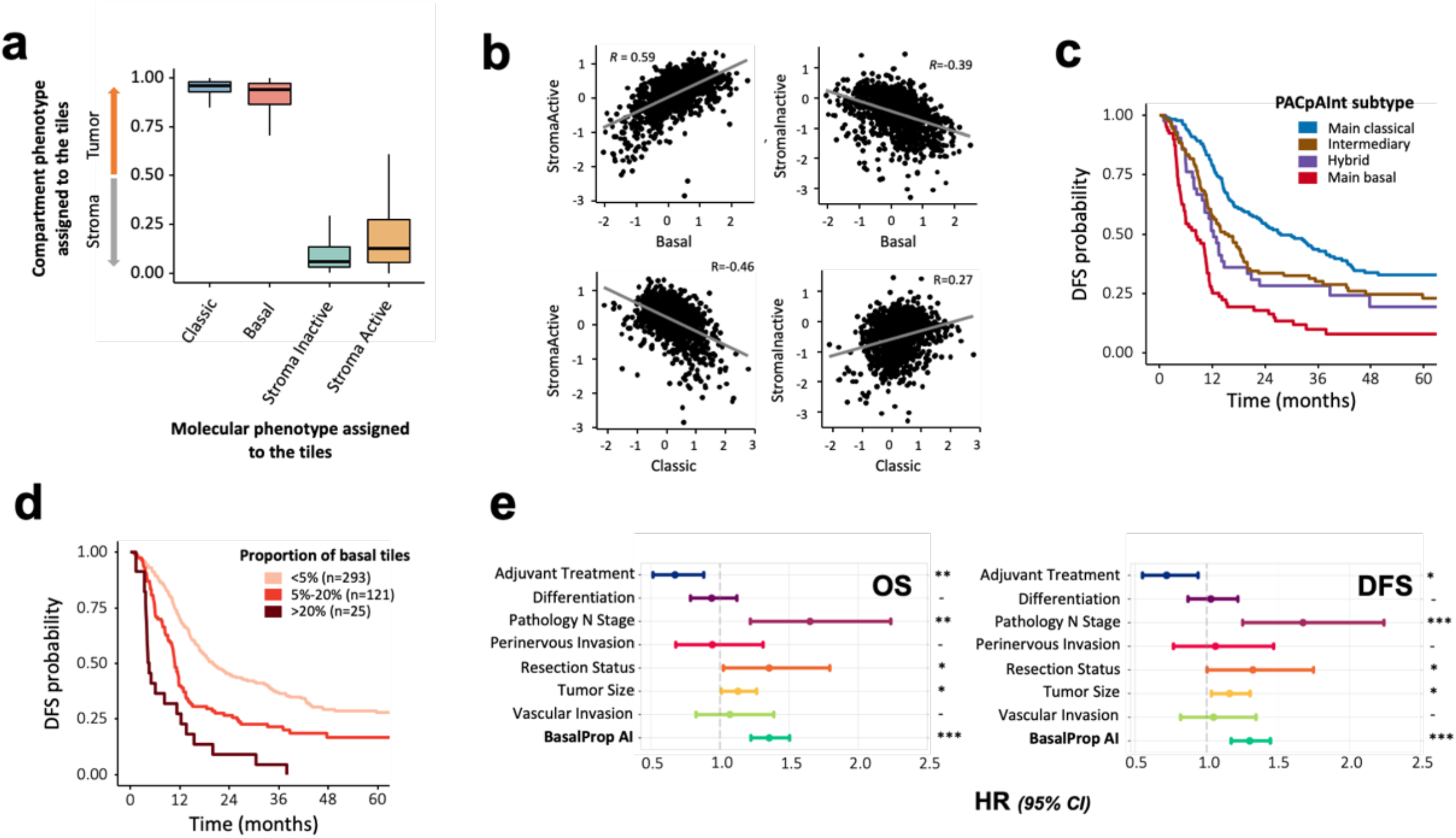
Identification of molecular components by PACpAInt-Comp. a) PACpAInt-Cell type tumor and stroma score in tiles identified as classical, basal-like, stroma active or inactive (analysis on 100K tiles), b) Correlation between slide-wise median stromal and epithelial scores, c) Kaplan Meyer analysis of disease free survival comparing pure classical, intermediate, hybrid and pure basal-like tumor, d) Stratification of patients using the proportion of basal-like tiles (less than 5%, 5% to 20%, more than 20%) and its association to disease-free survival, e) Multivariate analyses of clinical/pathological factors and PACpAInt-Comp computed amount of basal-like tile on overall (left) and disease free (right) survival.

